# Unraveling antiviral efficacy of multifunctional immunomodulatory triterpenoids against SARS-COV-2 targeting main protease and papain-like protease

**DOI:** 10.1101/2023.06.24.546363

**Authors:** Shweta Choudhary, Sanketkumar Nehul, Ankur Singh, Prasan Kumar Panda, Pravindra Kumar, Gaurav Kumar Sharma, Shailly Tomar

## Abstract

The coronavirus disease 2019 (COVID-19) pandemic, caused by severe acute respiratory syndrome coronavirus 2 (SARS-CoV-2) may be over, but its variants continue to emerge, and patients with mild symptoms having long COVID is still under investigation. SARS-CoV-2 infection leading to elevated cytokine levels and suppressed immune responses set off cytokine storm, fatal systemic inflammation, tissue damage, and multi-organ failure. Thus, drug molecules targeting the SARS-CoV-2 virus-specific proteins or capable of suppressing the host inflammatory responses to viral infection would provide an effective antiviral therapy against emerging variants of concern. Evolutionarily conserved papain-like protease (PLpro) and main protease (Mpro) play an indispensable role in the virus life cycle and immune evasion. Direct-acting antivirals targeting both these viral proteases represent an attractive antiviral strategy that is also expected to reduce viral inflammation. The present study has evaluated the antiviral and anti-inflammatory potential of natural triterpenoids: azadirachtin, withanolide_A, and isoginkgetin. These molecules inhibit the Mpro and PLpro proteolytic activities with half-maximal inhibitory concentrations (IC_50_) values ranging from 1.42 to 32.7 µM. Isothermal titration calorimetry (ITC) analysis validated the binding of these compounds to Mpro and PLpro. As expected, the two compounds, withanolide_A and azadirachtin, exhibit potent anti-SARS-CoV-2 activity in cell-based assays, with half-maximum effective concentration (EC_50_) values of 21.73 µM and 31.19 µM, respectively. The anti-inflammatory role of azadirachtin and withanolide_A when assessed using HEK293T cells were found to significantly reduce the levels of CXCL10, TNFα, IL6, and IL8 cytokines, which are elevated in severe cases of COVID-19. Interestingly, azadirachtin and withanolide_A were also found to rescue the decreased type-I interferon response (IFN-α1). The results of this study clearly highlight the role of triterpenoids as effective antiviral molecules that target SARS-CoV-2 specific enzymes and also host immune pathways involved in virus mediated inflammation.

## 1. INTRODUCTION

Despite the promising vaccination programs against coronavirus disease (COVID-19), vaccine hesitancy and compromised vaccine effectiveness has contributed to breakthrough infections, waning immunity, and the recurrence of new variants of severe acute respiratory syndrome coronavirus 2 (SARS-CoV-2). Whilst there have been tremendous strides forward in identifying drugs against SARS-CoV-2, no effective treatment for COVID-19 exists to date (1–3). One of the hallmark clinical feature of critically ill COVID-19 patients is the poor protective immune response with abnormally elevated levels of pro-inflammatory cytokines and chemokines such as interleukin-1 (IL-1), IL-2, IL-6, IL-8, IL-10, IL-12, IFN-γ, CXCL10, CXCL9, tumor necrosis factor-α (TNFα), etc. (3–6). Moreover, the induction of type I interferon (IFN-I; IFN-α1, IFN-β, IFN-δ, -ε, -κ, -τ) and type III interferon (IFN-III; IFN-λ1, -λ2, -λ3, and -λ4) responses are markedly impaired in severe COVID-19 patients (7–10). These dysregulated inflammatory responses lead to life-threatening pneumonia, multiorgan failure, acute respiratory distress syndrome (ARDS), and eventually death in more severe cases (2, 3, 6, 11). Since the host immune response also play a crucial role in disease progression, anti-inflammatory therapeutics capable of upregulating host antiviral response or inhibiting the systemic inflammation may prove to be a valuable option for clinical management of the disease (12).

SARS-CoV-2 has a remarkably large (∼30 kb), single-stranded RNA genome that encodes various structural, non-structural (nsP), and accessory proteins (13–15). Upon entry into the host cell, the genomic RNA is translated into pp1a and pp1ab polypeptides, which are processed by two virus-encoded cysteine proteases: the 3C-like protease (3CLpro or Mpro) and the papain-like protease (PLpro) (13–15). Subsequently, the processing of polyproteins releases 16 nsPs which compose the viral replication and transcription complex (RTC) that comprises RNA processing and proofreading function essential for maintaining the integrity of the genome (13, 15, 16). The multifunctional Mpro plays an essential role in processing two polypeptides pp1a and pp1ab, by operating at 11 sites, primarily at the conserved Leu-Gln↓(Ser/Ala/Gly) sequences (the cleavage site is indicated by ↓) (17–20). Active Mpro is a homodimer containing two protomers and each protomer is composed of a catalytic region comprising a noncanonical Cys-His dyad (21). In addition, Mpro targets the RIG-1 and the mitochondrial antiviral signaling (MAVS) protein–IFN signaling pathway to evade the innate immune responses of the host cell (22). Together with Mpro, PLpro also plays a vital role in the processing of viral polyprotein and the dysregulation of innate immune responses of the host. PLpro, a domain within nsP3, recognizes the P4-P1 consensus cleavage sequence LXGG found at the interface of nsP1/nsP2, nsP2/nsP3, and nsP3/nsP4 (23–25). This LXGG recognition sequence also resembles the sequence at the C-terminus of ubiquitin and ISG15 (23–25). Importantly, the deubiquitination and deISGylation of cellular proteins by PLpro are vital for antagonizing the antiviral pathways of the host (26, 27). The crystal structure of PLpro of SARS-CoV-2 signifies a right-handed architecture consisting of a ubiquitin-like (Ubl) domain and a “thumb-palm-finger” catalytic domain with a triad of Cys-His-Asp (24, 28). Due to the indispensable role of evolutionarily conserved Mpro and PLpro in the intracellular amplification stage of SARS-CoV-2, direct-acting antivirals against both of these proteins represent an attractive drug design strategy.

The unavailability of effective therapeutics has already exacerbated the severity of the COVID-19 pandemic. Plants have always been a rich repertoire of bioactive compounds with tremendous medicinal applications for centuries (29, 30). Compared with conventional drugs, phytopharmaceuticals from natural sources are considered a trending approach to cure ailments due to their affordability, efficacy, biocompatibility, bio-availability, and minimal side effects (31). Medicinal plants such as *Azadirachta indica*, *Withania somnifera*, *Tamarindus indica*, *Ocimum sanctum*, *Curcuma longa*, etc., are known to possess antiviral, antibacterial, or anti-cancerous activities due to the presence of phytochemicals such as flavonoids, terpenoids, limonoids, lignans, polyphenolics, coumarins, saponins, alkaloids, etc. (32–36). Of the numerous herbal plants, *Azadirachta indica* is a well-documented medicinal plant with virucidal action against human immunodeficiency virus (HIV), Chikungunya, Poliovirus, herpes simplex virus type-1 (HSV), Dengue, Chickenpox, etc. (37, 38). *Withania somnifera* also known as ’*Ashwagandha*’, is a prime herbal plant of Ayurvedic importance. Withanolides, a category of phytochemicals from *Withania somnifera,* have been reported to possess pharmacological activities such as an immunomodulation, prophylactic, antidepressant, anti-inflammatory, antioxidant, anti-tumor, and antiviral (39, 40). Ashwaganda’s immunomodulatory properties make it a potentially useful adjunct for treatment of viral infections including Human papillomavirus infection (HPV), Hepatitis C virus (HCV), HSV, and coronaviruses – including SARS-CoV and SARS-CoV-2 (41). *Ginkgo biloba* is an antioxidant-rich herb used as an anti-tumor, antibacterial, antiviral agent, and anti-inflammatory agent (42, 43). The most commonly reported bioflavonoid from *Ginkgo biloba,* isoginkgetin, exhibits antibacterial, anti-inflammatory, anti-tumor, and antiviral activities (43, 44). Considering the multifaceted role of plant-derived phytochemicals, identification of triterpenoids against virus-specific enzymes can be a promising approach to discover potent drug candidates for COVID-19 treatment.

In previous work, our group has identified three immunomodulatory triterpenoids (azadirachtin, withanolide_A, and isoginkgetin) as potential inhibitors of virus-encoded proteases (Mpro and PLpro) of SARS-CoV-2 using structure-assisted virtual screening and molecular dynamics approach (45, 46). Several *in silico* drug repurposing studies also predicted these triterpenoids to target Mpro and PLpro of SARS-CoV-2. However, former predictions were based solely on computational analysis and the efficacy data from their clinical trials against SARS-CoV-2 is missing (47–50). The present study investigates the antiviral and anti-inflammatory potential of identified triterpenoids in terms of Mpro and PLpro inhibition and cell-culture based SARS-CoV- 2 inhibition studies. The experimentally validated lead triterpenoids identified in this study present a dual-targeting virus-directed and host-directed strategy to avenge the pandemic of COVID-19. The findings in this study demonstrate the feasibility of this multi-targeting drug design strategy and pave the way for using triterpenoids to promote disease tolerance by suppressing immune-mediated pathology and impeding replication of SARS-CoV-2.

## 2. MATERIALS AND METHODS

### 2.1 Cells, viruses, and compounds

Vero cells (NCCS Pune, India) were maintained in high-glucose Dulbecco’s modified Eagle’s medium (DMEM; Himedia, India) complemented with 10% fetal bovine serum (FBS; Gibco, USA), 100 units of penicillin, and 100 µg streptomycin/mL (Himedia, India). DMEM augmented with 10% FBS, 100 units of penicillin, and 100 µg streptomycin/mL was used to culture HEK293T cells (NCCS Pune, India). All cells were grown at 37 °C in a humidified atmosphere with 5% CO_2_. SARS-CoV-2/Human/IND/CAD1339/2020 strain (GenBank accession no: MZ203529), isolated from SARS-CoV-2 infected Indian patient, was passaged in a culture flask of confluent Vero cells maintained in DMEM with 2% FBS (51). At the peak of infection, cell supernatant was harvested, centrifuged and further filtered through 0.22 µm filters to clarify cell debris. Virus stocks were kept at −80 °C until use. The virus was titered by end-point dilution method. All virus handling and experimental studies with live SARS-CoV-2 were conducted in Biosafety Level-3 facility at Indian Veterinary Research Institute (IVRI), Izatnagar, Bareilly, in accordance with the biosafety guidelines. Azadirachtin, withanolide_A, and isoginkgetin were procured from Cayman chemicals. Azadirachtin and withanolide_A were dissolved in methanol (Himedia, India) at 10 mM concentrations as stock solutions. Isoginkgetin was dissolved in Dimethyl sulfoxide (DMSO; Himedia, India) at 10 mM concentration.

### 2.2 Production of recombinant SARS-CoV-2 Mpro and PLpro

The plasmid encoding SARS-CoV-2 Mpro was a kind gift from Prof. Manidipa Banerjee, Indian Institute of Technology, Delhi. The gene corresponding to Mpro (1-306 amino acids) was amplified using 5’-ATATACCATGGTTAGCGGTTTTCGTAAAATGGCATTT-3’ as forward primer and 5’-TAATGCTCGAGCTGAAAGGTAACACCGCTACACTGAC-3’ as reverse primer. After restriction digestion, the gene was subcloned into *NcoI* and *XhoI* restriction sites in pET28c vector with an N-terminal histidine tag. Codon-optimized gene encoding the full-length SARS-CoV-2 PLpro (1-315 amino acids) was custom synthesized (Invitrogen, Thermo Fisher Scientific) and was cloned using *XhoI* and *NdeI* restriction sites in pET28c vector containing an N-terminal 6X histidine tag, using 5’-CAGCCATATGGAAGTTCGTACCATTAAAGT TTTTAC-3’ forward primer and 5’-GGTGCTCGAGCTATTTGATGGTGGTG-3’ reverse primer, as described previously (52). The recombinant constructs for PLpro and Mpro were transformed into competent *E. coli* Rosetta cells at 37 °C in Luria Broth (LB) containing 35 μg/mL chloramphenicol and 50 μg/mL of kanamycin until the optical density (O.D.) reached 0.6. The expression of Mpro and PLpro proteins were induced using 0.5 mM isopropyl-β-d-1- thiogalactopyranoside (IPTG) at 18 °C for ∼14 h. After centrifugation at 6000 rpm at 4 °C for 10 min, the PLpro and Mpro pelleted cells were resuspended in lysis buffer A (50 mM Tris-HCl, pH 8.5, 500 mM NaCl, and 5mM DTT) and lysis buffer B (50 mM Tris-HCl, pH 7.8, 150 mM NaCl), respectively. Next, the cells were lysed by French press (Constant Systems Ltd, Daventry, England), centrifuged at 10,000 rpm for 1 h, and the clarified supernatants were loaded onto Ni-NTA gravity flow column (BioRad) equilibrated with binding buffer for PLpro (50 mM Tris-HCl, pH 8.5, 500 mM NaCl, and 5mM DTT) and for Mpro (20 mM Tris-HCl, pH 7.8, 20 mM NaCl, and 5mM DTT). The columns were washed with binding buffer containing 5 mM imidazole and the target proteins were eluted with an elution buffer containing 100-500 mM imidazole. 12% SDS-PAGE was run to check the purity of the purified proteins (Supplementary figure S1a and S1c). Fractions containing pure PLpro and Mpro proteins were dialyzed against buffer C (50 mM Tris pH 8.5, 150 mM NaCl and 1 mM DTT) and buffer D (20 mM HEPES, pH 7.8, 20 mM NaCl, 1 mM DTT), respectively. The proteins were concentrated using 10 kDa cut off Amicon centrifugal filters (Millipore, USA).

### 2.3 Enzyme activity assay and IC_50_ determination

To determine the half-maximal inhibitory concentrations (IC_50_) of inhibitors against Mpro and PLpro, fluorescence-based cleavage assays were developed. Assay optimization for Mpro involved the usage of a fluorogenic peptide substrate Dabcyl-AVLQSGFR-Glu-Edans (Biolinkk, India), with Dabcyl as a quencher and Edans as a fluorophore, attached to the N- and C-terminals, respectively (Supplementary figure S1b). PLpro enzymatic activity was monitored using the fluorogenic substrate Z-RLRGG-7-amido-4-methylcoumarin (Biolinkk, India), with 7-amido-4- methylcoumarin as a fluorophore (Supplementary figure S1d). The proteolytic activity assays for PLpro and Mpro were performed in reaction buffer A (20 mM Tris-HCl pH 7.5, 150 mM NaCl) and reaction buffer B (20 mM Tris-HCl pH 7.8, 20 mM NaCl) respectively. The non-binding black 96-well microtiter plates (Corning) with a final reaction volume of 100 μL were used. The activity of SARS-CoV-2 was assessed by adding 1 μM purified enzymes to the plate followed by initiation of enzymatic reaction with the addition of 1.5 μM fluorogenic substrate peptides. The released fluorescence signals were monitored using a Synergy HT plate reader (BioTek Instruments, Inc) with wavelength filters for 360/40 nm (excitation) and 460/40 nm (emission). Inhibition of enzymatic activity of SARS-CoV-2 Mpro and PLpro were assayed by preincubating purified Mpro and PLpro proteins with compounds for 30 min at 25 °C. The substrate peptides were added in the last step at a final concentration of 1 μM/well. Upon the addition of substrate, the fluorescence signals were monitored at 360 nm (excitation) and 460 nm (emission). Measured fluorescence signals were blank-corrected with buffer containing different concentrations of the compounds incubated with the substrate without purified Mpro and PLpro enzymes. Data from duplicate sets of readings were plotted as dose inhibition curves to determine the IC_50_ values of compounds. Graph pad prism (version 8.3.1) and Microsoft Excel were used to analyze and prepare graphs.

### 2.4 Estimation of binding affinities by ITC

The dissociation constant and thermodynamic parameters for the affinity of compounds with proteases were determined using a MicroCal ITC200 micro calorimeter (Malvern, Northampton, MA). Binding of PLpro and Mpro with the compounds were measured in buffer A (50 mM Tris pH 8.5, 150 mM NaCl and 1 mM DTT) and buffer B (20 mM HEPES pH 8.0, 20 mM NaCl and 1 mM DTT), respectively. Titrations of Mpro were performed with 50 µM protein in the cell and 500 μM of inhibitors in the syringe at 25 °C. To test the binding between PLpro and compounds, 5 μM PLpro in the cell was titrated against 1 mM concentration of inhibitors in the syringe at 25 °C. For PLpro, a 0.5 μL priming injection followed by 13 consecutive 3 µl injections of compounds were titrated into 350 μL sample cell at 25 °C with 180 s intervals using a stirring rate of 750 rpm. For Mpro, a 0.5 μL priming injection followed by 15 consecutive 2.5 µl injections of compounds were titrated into 350 μL sample cell at 25 °C with 180 s intervals using a stirring rate of 750 rpm. All samples were gently vacuum-degassed before experiments. After subtracting the controls, the isotherm curves were fitted to a single-site binding model and the data was analysis was done using the Microcal Origin software provided by the manufacturer.

### 2.5 Cell viability assay

Cell viability profile of selected compounds was assayed in uninfected, compound-treated cells using MTT (3-(4, 5-dimethylthiazolyl-2)-2,5-diphenyltetrazolium bromide) assay. Monolayers of 1 × 10^4^ Vero cells/well in a 96-well plate were cultured in DMEM augmented with 10% FBS followed by an overnight incubation at 37 °C. On the next day, the media over cells was removed and fresh medium containing serially diluted compounds was added to each well of the plate followed by an incubation at 37 °C and 5% CO_2._ After 48 h of compound treatment, 20 μL of MTT reagent (5 mg/mL) (Himedia, India) was added to each well which was subsequently incubated for 4 h at 37 °C and 5% CO_2_ in the dark. Post incubation, the media was discarded and the formazan salt crystals were dissolved using 150 µL/well of DMSO. As readouts, absorbance values were measured at 550 nm using a Synergy HT plate reader (BioTek Instruments, Inc). Measurements from 0.1% DMSO-treated cells were included as positive control to calculate percent cytotoxicity for each compound. All treatments were made in triplicates. The CC_50_ (50% cytotoxic concentration) was calculated from the dose-response curve in GraphPad Prism 8.

### 2.6 Drug treatment and viral load reduction assay

To evaluate the antiviral spectrum of azadirachtin, withanolide_A, and isoginkgetin, Vero cells were seeded into 24-well cell-culture plate and were allowed to grow to confluency for 24 h at 37 °C and 5% CO_2_. Before virus inoculation, the cells were pre-treated with different doses of compounds for 3 h. Post incubation, the media containing compound was removed and the cells were washed with Phosphate Buffered Saline (PBS) (Himedia, India). SARS-CoV-2 at a multiplicity of infection (MOI) of 0.1 was added and incubated for 1.5 h. The virus inoculum was then removed and the cells were rewashed with PBS. Furthermore, fresh compound-containing DMEM media with 2% FBS was added. An equivalent volume of the methanol or DMSO solvent was added to virus control. At 48 h post-infection, the plate was freeze-thawed and the viral RNA in the cell lysate was extracted using HiPurA™ Viral RNA Purification Kit, according to the manufacturer’s instructions. The gene expression for Viral RNA in the cell lysate was quantified using quantitative real-time PCR (qRT-PCR) with the SARS-CoV-2-specific primers using the commercially available COVISure-COVID-19 Real-Time PCR kit (Genetix) according to the manufacturer’s protocol. Data was acquired using the following conditions: 15 min at 50 °C, a denaturation step at 95 °C for 2 min, 40 cycles of denaturation at 95 °C for of 30 s, and finally annealing at 60 °C for 30 s. All treatments were performed in duplicates for each compound. Percentage inhibition versus concentration graphs were plotted after normalizing the data using GraphPad Prism software to determine IC_50_ values.

### 2.7 Virus quantification assay

The production of infectious virus in compound treated cells and SARS-CoV-2 infected cells in comparison to only infected cells was quantified by tissue culture infectious dose 50 (TCID_50_) assay as reported previously (52). Vero cells were plated at a density of 1 × 10^4^ cells/well in a 96- well plate and the plate was incubated overnight at 37 °C in a 5% CO_2_ incubator to reach 80-90% confluency. The cellular monolayer was then exposed to 10-fold serial dilutions (10^1^ to 10^8^) of freeze-thawed lysate of cells infected and treated with compounds, collected at 48 h post-infection of the antiviral assay. Cells were incubated for 3 to 4 days at 37 °C in an incubator with 5% CO_2_. Cells were regularly observed for the appearance of obvious cytopathic effects (CPE). Mock-treated and infected samples were used as positive control, and samples with only cell-culture media were used as a negative control. After incubation, the viral inoculum was removed and the cells were fixed by the addition of 10% formaldehyde for 2 h followed by staining with 0.5% crystal violet solution. Virus endpoint titer was determined using the Reed-Muench method and expressed as log TCID_50_/mL.

### 2.8 Pro-inflammatory cytokine quantification assay

To ascertain the anti-inflammatory roles of selected triterpenoids, cell-based assays using HEK293T cells were optimized. In brief, HEK293T cells were seeded in 24-well plate at a density of 5.0 × 10^4^ cells per well in 10% DMEM supplemented 100 units of penicillin, and 100 µg streptomycin/mL and were incubated for 24 h for adherence. At a confluency of ∼80%, the cells were pre-treated with azadirachtin (75 µM) and withanolide_A (50 µM) diluted in OptiMEM media (Gibco). After an incubation period of 2 h, the compound containing media was removed and the cells were stimulated with 1 µg/mL of Polyinosinic:polycytidylic acid (poly I:C; Sigma) using Lipofectamine 2000 (Invitrogen) for 6 h. Poly I:C is a synthetic analog of dsRNA that mimics virus infection. Post incubation, the media over the cells was removed, fresh OptiMEM (Gibco) containing the compounds was added to cells, and the plate was incubated at 37 °C for 24 h. Untreated cells were included as controls. Total RNA was isolated from the cells using TRIzol (RNAiso; Takara) by following manufacturer’s instructions. In total, 1 µg RNA was subsequently converted to cDNA using random hexamer primer using PrimeScript™ 1st strand cDNA Synthesis Kit (Takara Bio) in accordance with manufacturer’s protocol. Gene expression analyses for IL6, IL-8, CXCL10, TNFα, and IFN-α1 cytokines were performed using KAPA SYBR fast universal qPCR kit (Sigma-Aldrich, USA) on QuantStudio real-time PCR system (Applied Biosystems, IN) following manufacturer’s instructions. Based on the ΔΔCt method, the relative expression of targeted genes was represented as fold changes with β-actin as endogenous control for normalization. All reactions were carried in duplicates in a 96-well plate (Applied Biosystems, IN). Fold change was calculated by taking mock-treated cells as 1.

The primer sequences used for qRT-PCR are as follows:

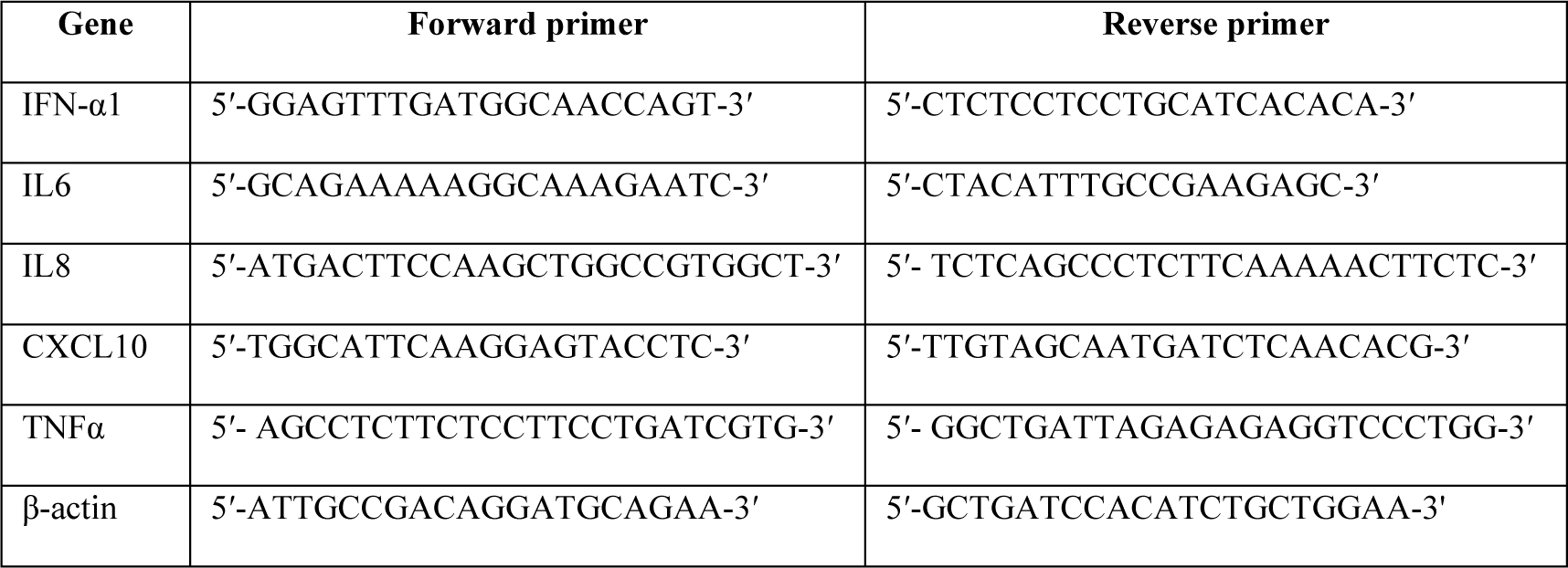

## 3. RESULTS

### 3.1 The potency of triterpenoids as a dual inhibitor against Mpro and PLpro

To dissect the inhibitory potential of the selected triterpenoids against SARS-CoV-2 proteases, fluorescence resonance energy transfer (FRET)-based assays were optimized for Mpro and PLpro. To accomplish this, gene sequences corresponding to Mpro and PLpro were cloned into the pET28c vector and were expressed in the *E. coli* expression system. Mpro and PLpro proteins were purified with Ni-NTA column to high purity (Supplementary figure S1a and S1c). To establish the FRET assay condition and to determine the inhibitory potency of identified compounds, fluorogenic substrate peptides corresponding to the cleavage sites for Mpro and PLpro were used. The selected triterpenoids displayed potent inhibitory activities against both the viral enzymes: the Mpro and PLpro, when tested in a concentration range of 0.1 to 100 µM. Dose-response experiments determined the IC_50_ values of azadirachtin, withanolide_A, and isoginkgetin by varying the compound concentrations against Mpro and PLpro of SARS-CoV-2. Withanolide_A, azadirachtin, and isoginkgetin inhibited the proteolytic activity of SARS-CoV-2 Mpro with IC_50_ values of 32.70 ± 3.20 µM, 19.85 ± 6.12 µM, and 21.34 ± 4.46 µM, respectively (Figure 1). For PLpro of SARS-CoV-2, the IC_50_ values 1.42 ± 0.57 µM, 15.56 ± 0.51 µM, and 7.5 ± 0.56 µM, were observed for withanolide_A, azadirachtin, and isoginkgetin respectively (Figure 1).

**FIGURE 1:**
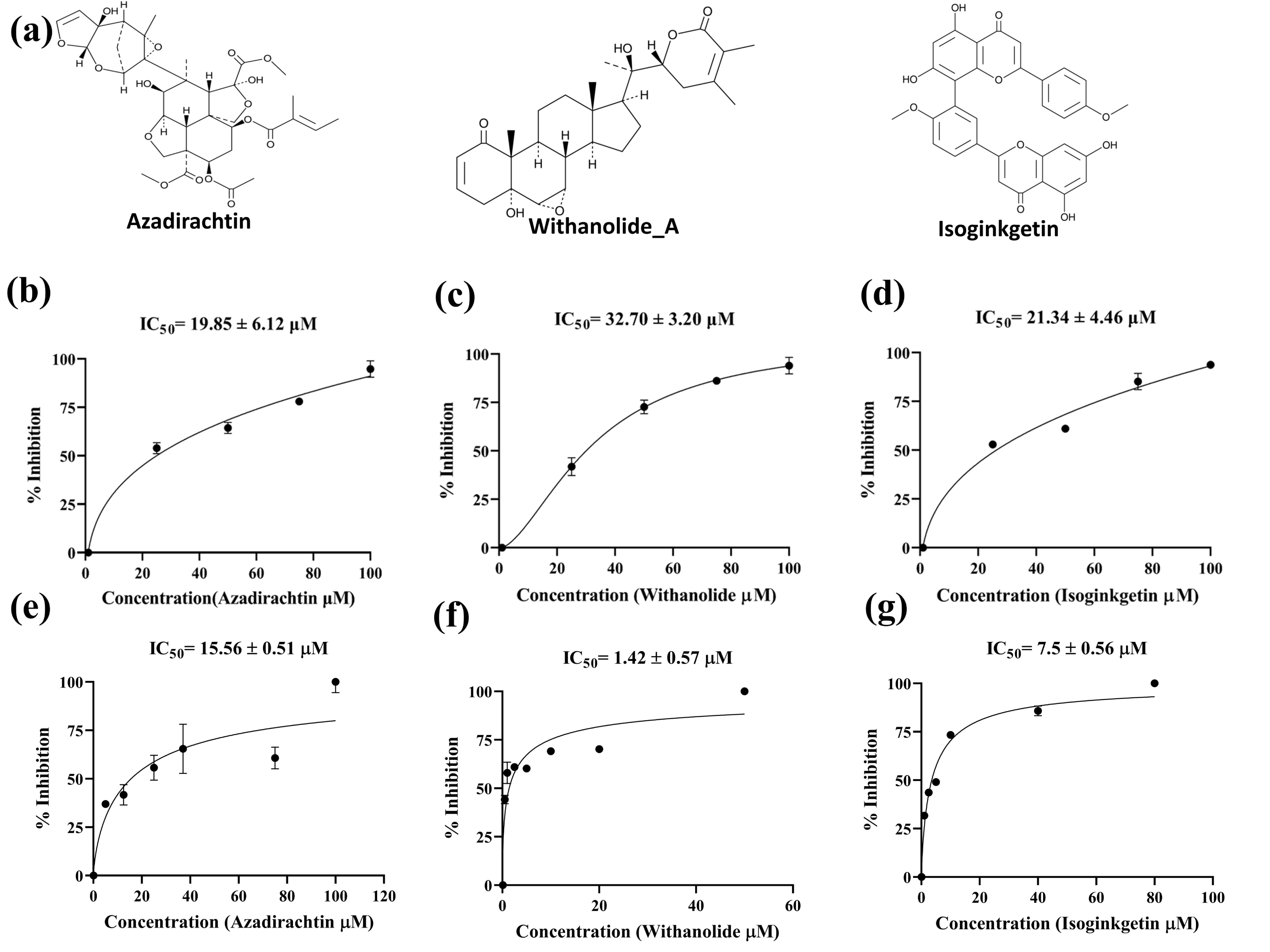
(a) Chemical structures of triterpenoids. (b, c, d) The inhibitory potencies of azadirachtin, withanolide_A, and isoginkgetin against the enzymatic activity of Mpro. (e, f, g) The inhibitory potencies of azadirachtin, withanolide_A, and isoginkgetin against the enzymatic activity of PLpro. Dose–response curves for IC_50_ values were determined by nonlinear regression fit. The calculated enzymatic activity with each compound was normalized to control. All data are shown as mean ± standard error, *n* = 2 biological replicates.

### 3.2 Estimation of binding affinities by ITC

To validate the binding of selected triterpenoids with SARS-CoV-2 PLpro and Mpro, the binding affinities with respective proteases were assessed using ITC. The thermodynamic data fitted into a one-site binding model revealed binding affinity (K_D_), of SARS-CoV-2 Mpro with withanolide_A, azadirachtin, and isoginkgetin as 127 µM, 88.4 µM, and 144 µM, respectively (Table 1 and figure 2). With regard to SARS-CoV-2 PLpro, the ITC experiments revealed K_D_ values of 478 µΜ, 641 µΜ, and 169 µΜ for withanolide_A, azadirachtin, and isoginkgetin, respectively. The results of thermodynamic parameters are documented in Table 1. Moreover, the 1:1 stoichiometry (N) detected postulates specificity for a single binding site of Mpro and PLpro (Table 1). The binding of withanolide_A, azadirachtin, and isoginkgetin to SARS-CoV-2 PLpro and Mpro is exothermically driven as shown by their enthalpy variation (Δ*H*) (Table 1 and figure 2). As depicted in Figure 2 and Table 1, the measured thermodynamic parameters and the titration curves revealed good binding affinities of Mpro and PLpro with the selected triterpenoids, which nicely corroborates with the data from FRET assay.

**FIGURE 2:**
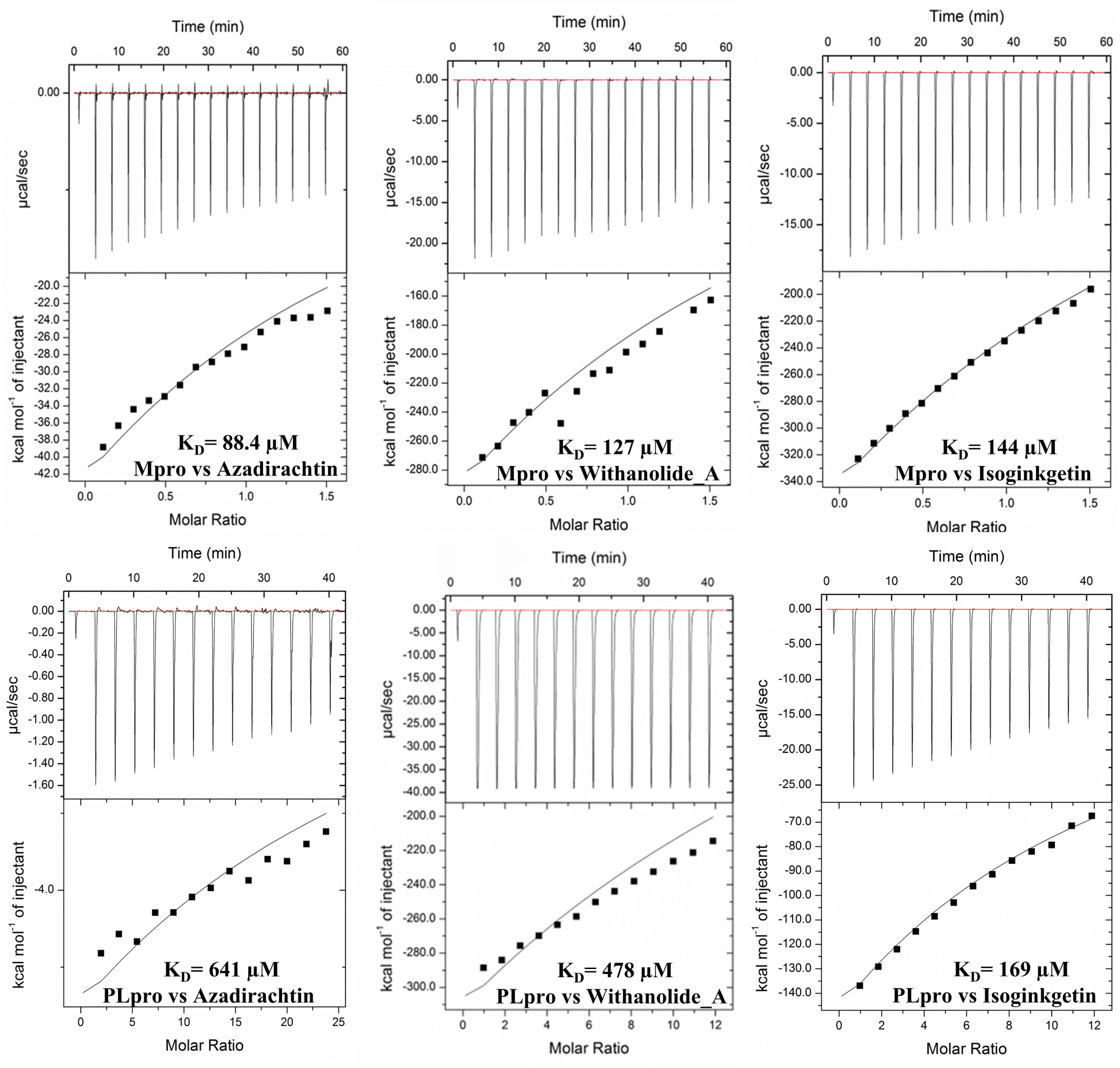
Binding analysis of azadirachtin, withanolide_A, and isoginkgetin with SARS-CoV-2 Mpro and PLpro characterized by ITC. The top of each ITC titration is its raw data and the bottom represents the binding isotherm fitted to one sets of site model. K_D_ values are inset.

**Table 1:** Thermodynamic binding parameters of azadirachtin, withanolide_A, and isoginkgetin for purified PLpro and Mpro of SARS-CoV-2.

| Compound | No. of sites (N) | K<sub>D</sub> (μM) | ΔH (cal/mol) | ΔS (cal/mol/degree) |
| --- | --- | --- | --- | --- |
| **PLpro** |  |  |  |  |
| Withanolide_A | 1 | 478 | $-2.952 \times 10^7 \pm 5.136 \times 10^6$ | $-9.90 \times 10^4$ |
| Azadirachtin | 1 | 641 | $-2.927 \times 10^5 \pm 1.1$ | $-9.67 \times 10^2$ |
| Isoginkgetin | 1 | 169 | $-4.949 \times 10^6 \pm 1.55$ | $-1.66 \times 10^4$ |
| **Mpro** |  |  |  |  |
| Withanolide_A | 1 | 127 | $-2.47 \times 10^6 \pm 9.031 \times 10^4$ | $-8.27 \times 10^3$ |
| Azadirachtin | 1 | 88.4 | $-1.631 \times 10^5 \pm 1.89 \times 10^4$ | $-5.29 \times 10^2$ |
| Isoginkgetin | 1 | 144 | $-1.641 \times 10^6 \pm 2.809 \times 10^5$ | $-5.49 \times 10^3$ |

N: binding stoichiometry; K<sub>D</sub>: equilibrium dissociation constant; ΔH: enthalpy variation; ΔS: entropy variation; SD: standard-deviation;

### 3.3 Triterpenoids inhibit SARS-CoV-2 *in vitro*

To further authenticate the enzyme inhibition and ITC results, the capability of the selected compounds to inhibit SARS-CoV-2 *in vitro* was evaluated. Among the tested inhibitors, withanolide_A and azadirachtin displayed no cytotoxicity at tested concentrations (Figure 3). The CC_50_ values of both withanolide_A and azadirachtin are above 100 µM, indicating a good safety window for these triterpenoids. However, there was an observed increase in cellular cytotoxicity in the presence of the compound isoginkgetin at concentrations of more than 1 µM (Supplementary figure S2). To determine the effectiveness of triterpenoids on SARS-CoV-2, Vero cells were treated with withanolide_A, azadirachtin, and isoginkgetin and viral infection was quantified by qRT-PCR. Percent inhibition data were normalized to the infected controls (solvent DMSO or methanol treated as well as infected cells) and cell-only control (Solvent DMSO or methanol treated). The percent cytotoxicity data was normalized to cell only control (solvent DMSO or methanol treated). In line with previous observations, cell-based assays revealed that treatment with withanolide_A and azadirachtin resulted in significant inhibitory potential against SARS-CoV-2 with a half-maximum effective concentration (EC_50_) value of 21.73 µM and 31.19 µM, respectively. Furthermore, both compounds harbored discernable dose-dependent antiviral activities with a percentage inhibition of more than 90% (Figure 3a and 3b). However, the compound isoginkgetin was not observed to inhibit the replication of SARS-CoV- 2 at its non-toxic concentrations (Supplementary figure S2). Consistent with these viability assay results, the high cytotoxicity for isoginkgetin observed in this study is in accordance with other published studies (43).

**FIGURE 3:**
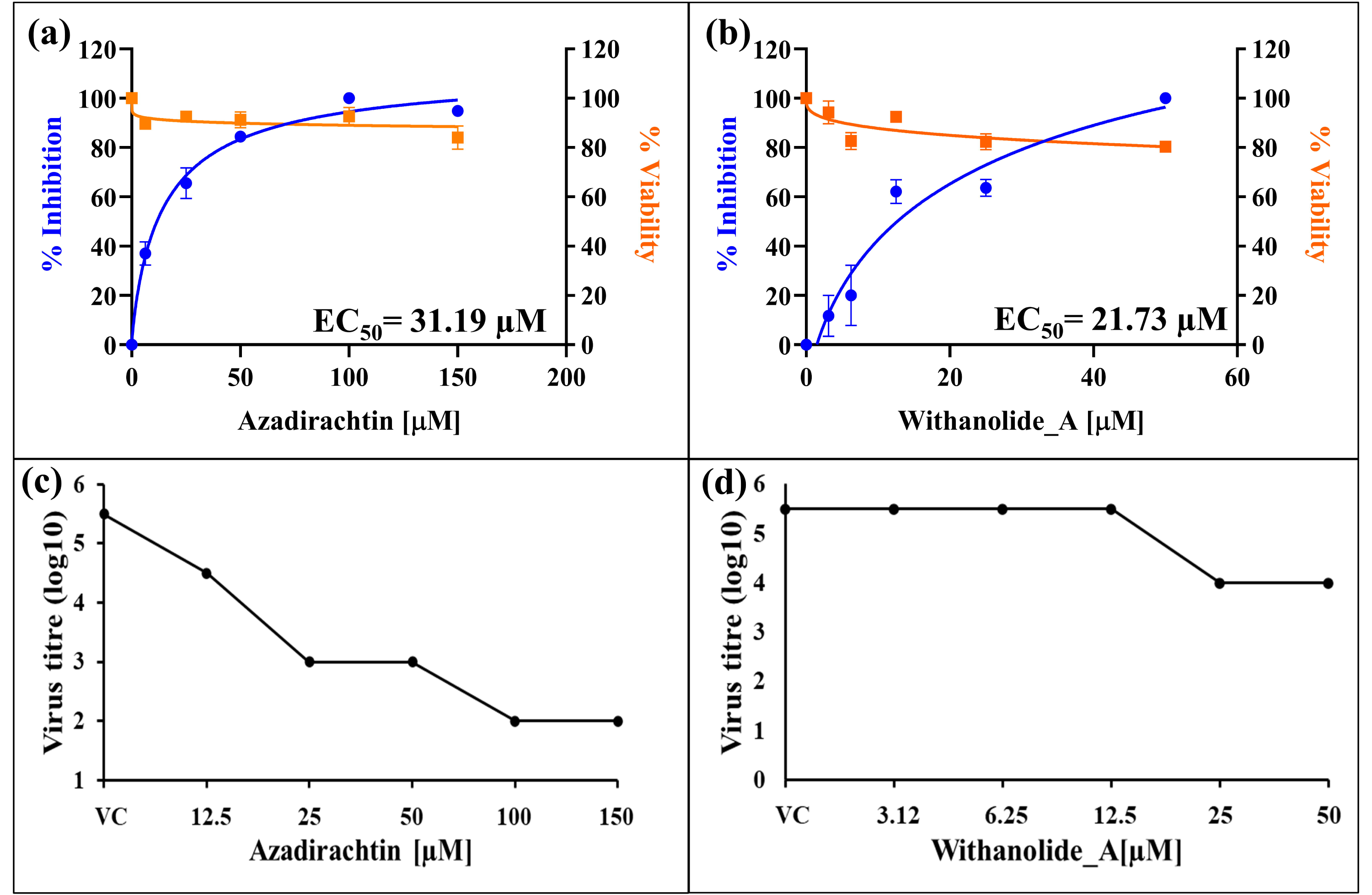
Immunomodulatory triterpenoids reduce SARS-CoV-2 infection. (a and b) Vero cells were infected with SARS-CoV-2 (0.1 MOI) in the presence of indicated concentrations of compounds for 48 h followed by the quantification of viral RNA copies by qRT-PCR and TCID_50_ assay. Cytotoxicity of compounds on Vero cells was measured using the MTT assay. MTT assay data represents mean and standard deviations from triplicates (n=3). Percent inhibition of SARS-CoV-2 by azadirachtin and withanolide_A on Vero cells (represented in blue) and percent viability of azadirchtin and withanolide_A in Vero cells (represented in orange). Percentage inhibition values are represented as mean ± SD of two independent experiments. Error bars corresponds to the standard deviation from duplicate experiments. (c and d) SARS-CoV-2 viral titer after treatment with triterpenoids. Confluent monolayer of Vero cells in a 96-well plate were inoculated with 10-fold serial dilutions of samples harvested from virus yield reduction assay and the plate was incubated for 3-4 days until the appearance of cytopathic effect (CPE). After its incubation period, 10% formaldehyde was used to fix the cells and staining was done using crystal violet solution. Data is expressed as log TCID_50_/mL.

Encouraged by the promising antiviral activity of withanolide_A and azadirachtin, a further set of experiments were performed to determine the TCID_50_ for azadirachtin and withanolide_A. Treatment with azadirachtin reduced virus titer from 5.5 log TCID_50_/mL to 2 log TCID_50_/mL (Figure 3c). Withanolide_A decreased the viral titer to 4 log TCID_50_/mL compared to mock-treated virus control (5.5 log TCID_50_/mL) (Figure 3d). Indeed, a more significant decrease in infectious viral progenies was observed when virus-infected cells were treated with higher concentrations of azadirachtin fold (3.5 log reduction TCID_50_/mL) in comparison to withanolide_A (decrease of 1.5 log TCID_50_/mL) (Figure 3c and 3d). The results obtained from the TCID_50_ assay demonstrated a dose-dependent decrease in infectious viral titer. Collectively, the data provides strong evidence that azadirachtin and withanolide_A could potentially impede SARS-CoV-2 replication *in vitro*.

### 3.4 Anti-inflammatory role of azadirachtin and withanolide_A

SARS-CoV-2 infections are often associated with highly elevated levels of pro-inflammatory cytokines, including IL-6, IL-1, IL-8, IL-10, IL-12, TNFα, IFN-γ, CXCL10, and CXCL9 that in many cases lead to severe lung damage and multiorgan failure (2–5, 11, 53). Previous studies have also reported lymphopenia, neutrophilia, reduced or delayed type I interferon responses (IFNα and IFNβ) and hyperproduction of pro-inflammatory cytokines, which are primarily responsible for aggravating immunopathological complications in COVID-19 patients (7–10). Therefore, an immunomodulatory therapy to control the virus-triggered exuberant cytokine release can provide wide-spectrum effect in controlling SARS-CoV-2 infection and resuming host’s homeostasis. The expression of inflammatory mediators stimulated on poly I:C treatment was therefore examined to see if azadirachtin and withanolide_A had any anti-inflammatory effects. Cellular lysates collected after 24 h of poly I:C stimulation were quantified for the levels of cytokine production (CXCL10, TNFα, IL6, IL8, and IFN-α1) using qRT-PCR. The stimulation of HEK293T cells with Poly I:C dramatically increased the levels of pro-inflammatory cytokines CXCL10, TNFα, IL6, and IL8 when compared with untreated cellular control (Figure 4). Interestingly, the mRNA expression levels of CXCL10, TNFα, IL6, and IL8 upregulated in response to poly I:C were observably decreased after treatment with azadirachtin and withanolide_A (Figure 4). Further analysis also showed that treatment with azadirachtin and withanolide_A also restored the levels of IFN-α1, that are reported to be remarkably reduced in COVID-19 patients (Figure 4) (3, 7). Altogether, the data from SARS-CoV-2 antiviral assays and anti-inflammatory studies speculate that these triterpenoids inhibit both viral replication as well as the expression of inflammatory genes.

**FIGURE 4:**
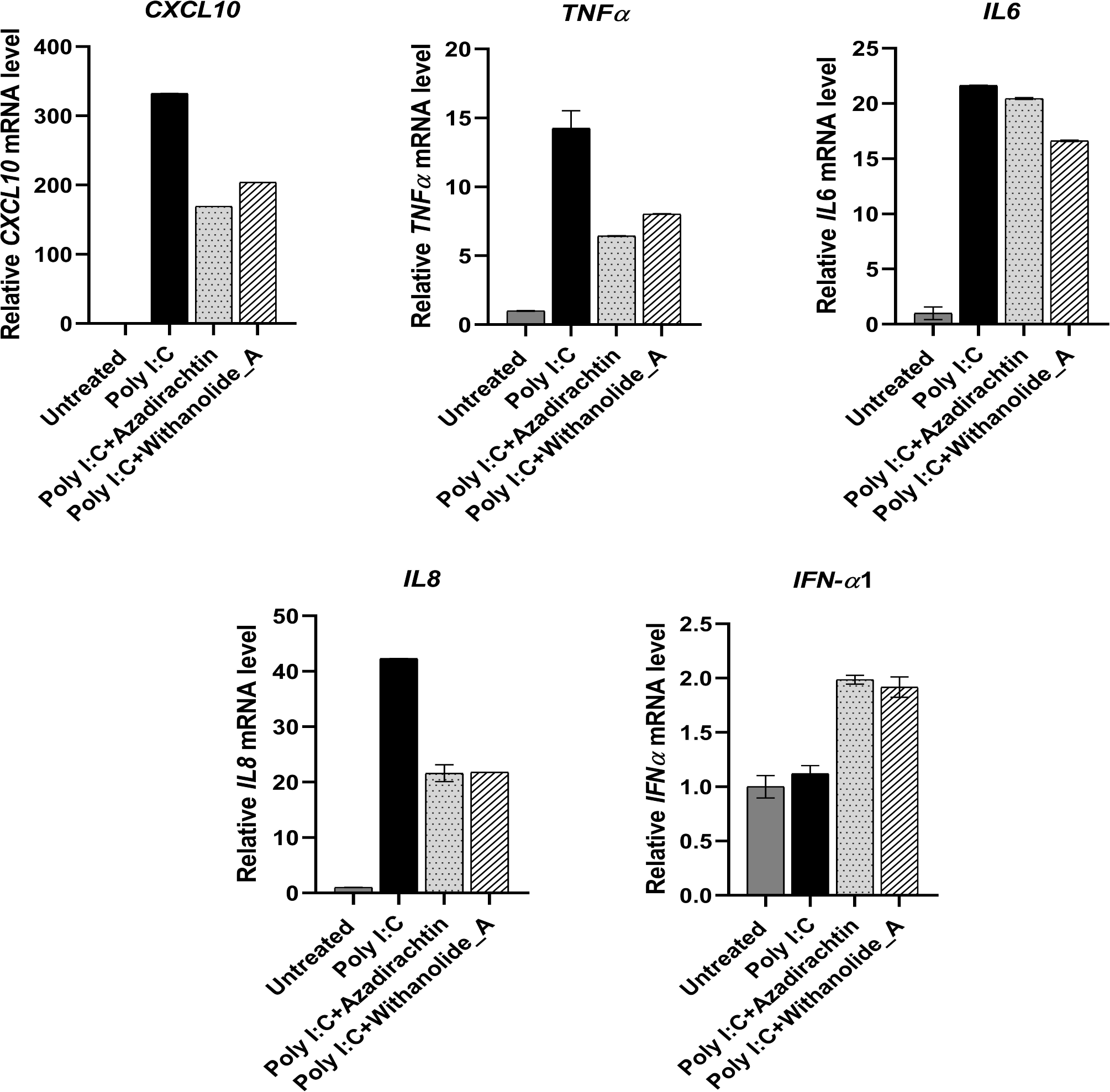
Anti-inflammatory effects of azadirachtin and withanolide_A. HEK293T cells pre-treated with azadirachtin (75 µM) and withanolide_A (50 µM) for 2h, followed by stimulation with Poly I:C (1 µg/mL) for 6 h. Post incubation, the cells were allowed to grow in fresh medium containing compounds. 24 h post poly I:C treatment, the total RNA was extracted and the relative fold change of CXCL10, TNFα, IL6, IL8, and IFN-α1 was determined using qRT-PCR. Expression of each gene was normalized to endogenous control β-actin and gene expression for untreated cells was taken as a reference. Shown are the data for mean ± SD of duplicate reactions.

## 4. DISCUSSION

In the setting of waning immunity from antibody-based interventions for SARS-CoV-2, broadly effective antiviral therapies with multiple modes of action have increased in importance and might become an important tool in fight against the COVID-19 pandemic. Despite advances in modern vaccine and therapeutics, the landscape of effective treatment options against SARS-CoV-2 is still underdeveloped. So far, the U.S. Food and Drug Administration (FDA) approved treatment options against SARS-CoV-2 includes remdesivir, molnupiravir, and paxlovid, which are reported to shorten COVID-19 hospitalization times, but exhibited limited efficacy in clinical use (54–57). The requirement for effective antiviral therapies is still a crucial aspect for timely management of viral infection and to improve the preparedness for future outbreaks of SARS-CoV-2 or its other emerging variants (58, 59). A strategy combining virus-specific and host-targeted antivirals can potentially offer effective antiviral therapies to combat emerging variants of SARS-CoV-2 and to reduce the virus induced inflammatory responses in mild to severely infected COVID-19 patients (60, 61). SARS-CoV-2 specific proteases, the Mpro and PLpro, are of particular interest for developing effective antiviral therapies because of their fundamental roles in viral replication and evasion of innate immune responses (18, 19, 23, 62–64). Additionally, the conservation of substrate-binding pockets in both proteases across coronaviruses is expected to provide effective therapeutic interventions to treat COVID-19 amid the emergence of new variants (64).

Of importance, medicinal plants offer a rich reserve of phytochemicals with immense therapeutic potentiality as antimicrobial, antiviral, anti-tumor, anti-inflammatory, neuroprotective, cardioprotective, and immunomodulatory effects (29, 30, 32, 33, 65). The triterpenoids are an important class of secondary metabolites from natural sources with wide-spectrum pharmacological activities including antiviral, antibacterial, anti-tumor, anti-cancer, anti- microbials, etc. (30, 42, 43, 49, 66–70) Triterpenoids have also been found to play immunomodulatory or anti-inflammatory roles highlighting their potential in management of immunological complications (32, 39, 40, 44, 49, 71). In this study, the three triterpenoids (azadirachtin, withanolide_A, and isoginkgetin) which were earlier identified by *in-silico* analysis targeting the SARS-CoV-2 virus specific proteases (Mpro and PLpro), have been further investigated for their antiviral as well as anti-inflammatory roles (45, 46). Given that the triterpenoids target the active site of both proteases (45, 46), the direct inhibitory effects on enzymatic activities of Mpro and PLpro were measured using an *in vitro* fluorescence-based protease assay. The selected compounds displayed multi-potent inhibitory effects against proteolytic activities of both Mpro and PLpro of SARS-CoV-2 with IC_50_ values in the low micromolar range (Figure 1).

To investigate the binding affinities and stoichiometry of inhibitors to Mpro and PLpro, ITC experiments were conducted that confirmed the stoichiometry of the compounds for a single binding site (Figure 2). The data from the ITC experiment revealed good binding affinities of Mpro with the K_D_ as 88.4 μM, 127 μM, and 144 μM, for azadirachtin, withanolide_A, and Isoginkgetin, respectively (Figure 2, Table 1), which nicely corroborates with the result of Mpro proteolytic assay. The binding affinities of PLpro with withanolide_A, azadirachtin, and isoginkgetin was also examined that revealed K_D_ values of 478 µΜ, 641 µΜ, and 169 µΜ, respectively (Figure 2 and Table 1). Treatment with azadirachtin and withanolide_A exhibited robust antiviral effects in *in vitro* cell-based assays against SARS-CoV-2 with EC_50_ values of 31.19 µM and 21.73 µM, respectively (Figure 3). Hemdan *et al*. proposed the antiviral activity of extracts from leaves and bark of *Azadirachta indica* against SARS-CoV-2, however the exact mechanism of action was not provided (66). The present study identified azadirachtin, an important triterpenoid from *Azadirachta indica,* to be one of the bioactive molecules responsible for exhibiting inhibitory activity against SARS-CoV-2 and its proteases. Consistent with previously published reports, the compound isoginkgetin displayed considerably higher cellular toxicity (43), and was not observed to inhibit viral replication in Vero cells in this study at concentrations lower than those exhibiting cytotoxicity (Supplementary figure S2).

Infection with SARS-CoV-2 can lead to an overexuberant immune response that can lead to severe pneumonia, cytokine storm, multiorgan dysfunction, and eventually death (2–4, 11). A dysregulated immune response associated with attenuated IFN responses and overproduction of inflammatory cytokines or chemokines (CXCL10, TNFα, IL6, IL8) has been proposed to be a prime reason for cytokine storm or ARDS (1–3, 7). Therapies focused on modulating innate immune responses to attenuate the excessive release of pro-inflammatory cytokines or to promote the IFN antiviral response represent a noteworthy target for treating patients with severe COVID- 19 (72, 73). Published literature suggests a wide-spectrum of pharmacological properties of both azadirachtin and withanolide_A such as anticancer, anti-inflammatory, antiviral, antibacterial, and immunomodulators (32, 33, 39, 40, 68, 69). Owing to the significant inhibitory effects against SARS-CoV-2 and safer toxicity profiles, only azadirachtin and withanolide_A were selected for further studies. To ascertain the roles of withanolide_A and azadirachtin on virus-triggered inflammatory pathways, HEK293T cells were treated with poly I:C, a TLR3 agonist, that mimics virus infection and induces inflammatory responses in the host cells (67, 74, 75). Findings in this study revealed that treatment with azadirachtin and withanolide_A decreased the levels of CXCL10, TNFα, IL6, IL8 that are well documented to be modulated in critically ill COVID-19 patients (Figure 4). Previous studies have shown the regulatory roles of SARS-CoV-2 Mpro and PLpro in delayed type I IFN responses as an immune evasion strategy (22, 27, 64, 76). Interestingly, both compounds were able to rescue the decreased IFN-α1 responses, which could possibly be due to inhibition of both Mpro or PLpro or any other off-target in signaling cascade (Figure 4). Altogether, the present study supports the notion that triterpenoids effectively inhibit the enzymatic activity of Mpro and PLpro, reduce SARS-CoV-2 viral load, and reduces inflammatory signatures.

Considering the dual inhibitory effect of withanolide_A and azadirachtin on both viral replication and inflammatory markers, the findings from this study evidently suggests that the triterpenoids could efficiently inhibit the replication of SARS-CoV-2 and ameliorate cytokine storm induced immunological injuries. The favorable profiles and SARS-CoV-2 antiviral activity of withanolide_A and azadirachtin supports further *in vivo* and clinical testing for the treatment of COVID-19.

## AUTHOR CONTRIBUTIONS

ST, PK, PKP, and GKS planned and designed the study together. ST, PK, and GKS supervised the experimental work. ST, PK, PKP, and GKS analyzed and discussed the results. SC, AS, and SN performed the experiments, analyzed the data with critical inputs from ST, PK, GKS, and PKP. The manuscript was prepared by ST, PK, GKS, SC, AS, PKP, and SN. All authors read and approved the final manuscript.

## Supporting information

supplementary file updated

## ACKNOWLEDGMENTS

This work was supported by funding from Science and Engineering Research Board, Department of Science & Technology, Government of India (Project reference no. IPA/2020/000054). The authors acknowledge and thank Bioinformatics Center (BIC) supported by Department of Biotechnology, Govt of India (reference number BT/PR40141/BTIS/137/16/2021), Ashok Soota Molecular medicine facility, and the Macromolecular Crystallographic Unit (MCU) at Indian Institute of Technology, Roorkee. We thank Prof. Manidipa Banerjee for providing Mpro plasmid. SC thank Council of Scientific & Industrial Research, Government of India for financial support. AS and SN acknowledges the Ministry of Human Resource Development, (MHRD) for financial support.

