## supplementary file updated for "Unraveling antiviral efficacy of multifunctional immunomodulatory triterpenoids against SARS-COV-2 targeting main protease and papain-like protease"

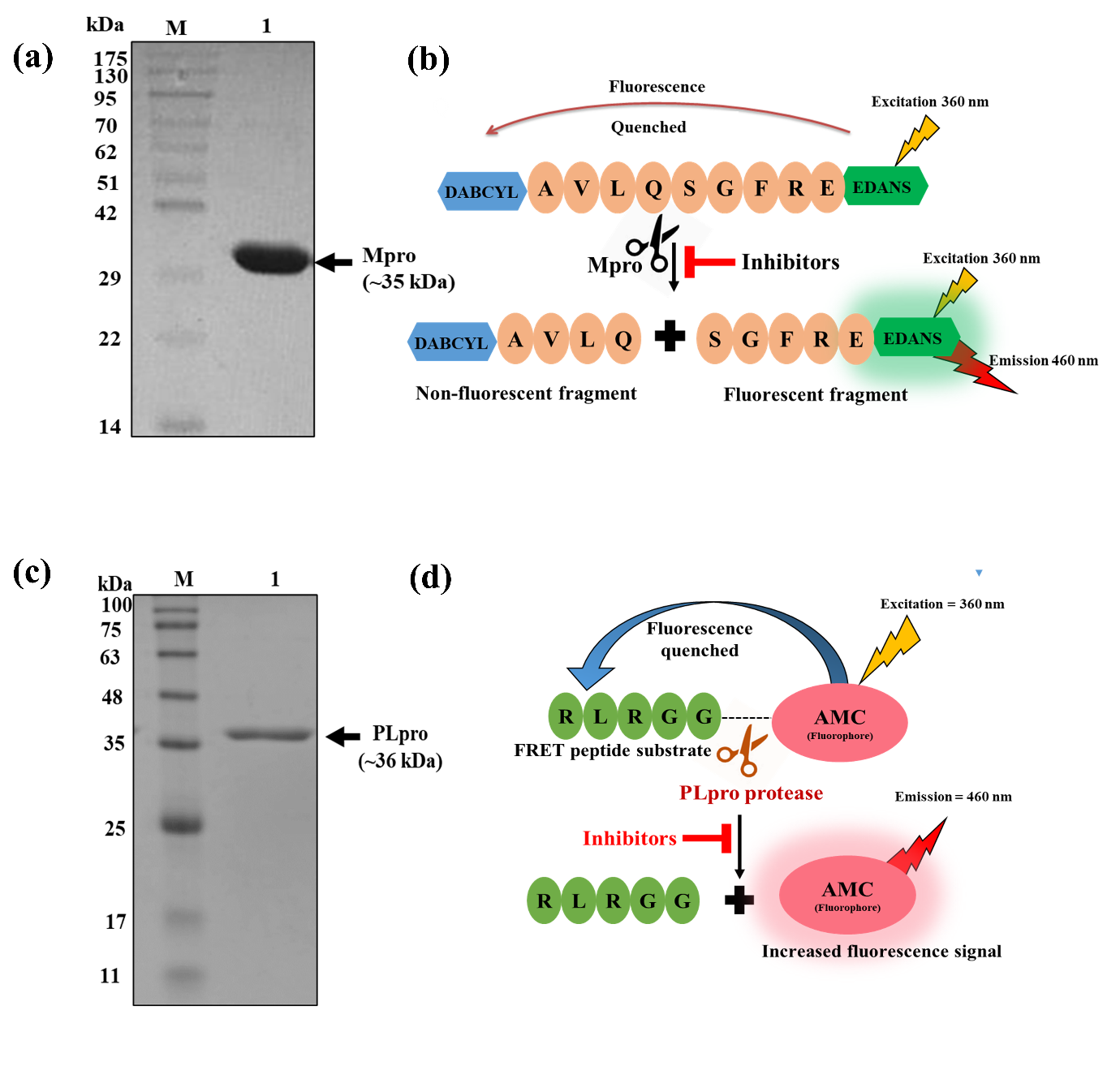

**Supplementary figure S1:** **Purification and characterization of SARS-CoV-2 Mpro and PLpro.** (a) SDS-PAGE of purified Mpro. Lane M: protein ladder; lane 1: Histidine tagged-Mpro. (b) Development of high-throughput FRET based enzymatic assay for Mpro. A fluorogenic FRET substrate Dabcyl-AVLQSGFR-Glu-Edans, with Edans as a fluorophore (shown in green) and Dabcyl as a quencher (shown in blue) is represented. Mpro cleaves the peptide sequence and separate the fluorescent molecule from the quencher, which results in increase in the fluorescence signal. In presence of inhibitors in the reaction mixture, the fluorescence signal will get reduced. (c) SDS-PAGE of purified PLpro. Lane M: protein ladder; lane 1: Histidine tagged-PLpro (d) Schematic for fluorogenic substrate Z-RLRGG-7-amido-4-methylcoumarin for PLpro, with 7-amido-4-methylcoumarin (AMC) as a fluorophore. Release of the C-terminal AMC dye by proteolysis activity of PLpro generates a fluorescence signal. The intensity of signal gets decreased in the presence of PLpro inhibitors.

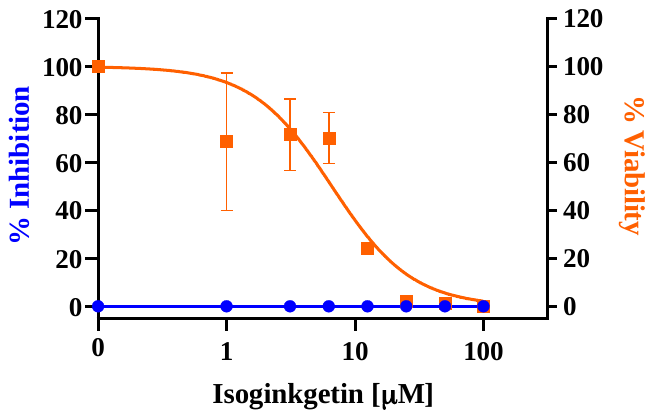

**Supplementary figure S2:** Immunomodulatory triterpenoids reduce SARS-CoV-2 infection. Vero cells were infected with SARS-CoV-2 (0.1 MOI) in the presence of indicated concentrations of isoginkgetin for 48 h followed by the quantification of viral RNA copies by qRT-PCR and TCID_50_ assay. Cytotoxicity of isoginkgetin on Vero cells was measured using the MTT assay. Percent inhibiton of SARS-CoV-2 by isoginkgetin on Vero cells is represented in blue and percent viability of isoginkgetin in Vero cells is represented in orange. Values are represented as mean ± SD of two independent experiments. Error bars represent the standard deviation from duplicate experiments.
